# Honey bees can store and retrieve independent memory traces after complex experiences that combine appetitive and aversive associations

**DOI:** 10.1101/2021.10.12.464105

**Authors:** Martín Klappenbach, Agustín E. Lara, Fernando F. Locatelli

**Affiliations:** Departamento de Fisiología, Biología Molecular y Celular, Facultad de Ciencias Exactas y Naturales, Instituto de Fisiología, Biología Molecular y Neurociencias, Universidad de Buenos Aires-CONICET), Ciudad Universitaria, Buenos Aires, Argentina

## Abstract

Real-world experiences do often mix appetitive and aversive events. Understanding the ability of animals to extract, store and use this information is an important issue in neurobiology. We used honey bees as model to study learning and memory after a differential conditioning that combines appetitive and aversive training trials. First of all, we describe an aversive conditioning paradigm that constitutes a clear opposite of the well known appetitive olfactory conditioning of the proboscis extension response. A neutral odour is presented paired with the bitter substance quinine. Aversive memory is evidenced later as an odour-specific impairment in appetitive conditioning. Then we tested the effect of mixing appetitive and aversive conditioning trials distributed along the same training session. Differential conditioning protocols like this were used before to study the ability to discriminate odours, however they were not focused on whether appetitive and aversive memories are formed. We found that after a differential conditioning, honey bees establish independent appetitive and aversive memories that do not interfere with each other during acquisition or storage. Finally, we moved the question forward to retrieval and memory expression to evaluate what happens when appetitive and the aversive learned odours are mixed during test. Interestingly, opposite memories compete in a way that they do not cancel each other out. Honey bees showed the ability to switch from expressing appetitive to aversive memory depending on their satiation level.

## Introduction

Animals gather and integrate information from the environment to take appropriate decisions (Tinbergen, 1951). In this process, the value of a detected signal must be computed in the context of concurrent cues that may reinforce the meaning of the first one or contradict it and demand different actions (Lewis et al., 2015). Furthermore, the relevance of a signal might change depending on the internal state of the animal. For example, an animal that has recently eaten looses interest in food signals (Yapici, Cohn, Schusterreiter, Ruta, & Vosshall, 2016), or the effort to find a mate following a pheromone plume vanishes after mating (Zhang, Rogulja, & Crickmore, 2016). Thus, stimuli that are highly important at a given moment can be irrelevant minutes later. Interestingly, the need of these computations does not only apply to signals with innate meaning but also to learnt cues. How all these external and internal stimuli are integrated to drive adaptive decisions constitutes fundamental questions in neurobiology (Davis, 1979; Sugrue, Corrado, & Newsome, 2005). In this context, animal models and behavioural learning paradigms that allow fine control of external and internal stimuli are instrumental to disentangle these computations. Classic experimental approaches to study learning and memory take benefit of experimental designs that involve single appetitive or aversive associations. In these cases, a conditioned stimulus or action is associated with a reward or punishment, and therefore the changes in behaviour can be linked to this single association. Thanks to such clear approaches, the instructive role that distinct neural pathways and biogenic amines play in mediating aversive and appetitive learning was successfully described in several invertebrate models (Aso & Rubin, 2016; Burke et al., 2012; Farooqui, Robinson, Vaessin, & Smith, 2003; Kaczer & Maldonado, 2009; Klappenbach, Maldonado, Locatelli, & Kaczer, 2012; Mizunami & Matsumoto, 2017; Totani et al., 2019; Vergoz, Roussel, Sandoz, & Giurfa, 2007). Since that, a research field of growing interest has been to understand how appetitive and aversive stimuli that often appear mixed in realistic experiences, interact during learning and memory retrieval to coordinate adaptive decisions and behaviour. With this idea in mind, several works started studying learning and memory in situations in which stimuli and memories of opposite valence compete during learning or retrieval (Das et al., 2014; Felsenberg et al., 2018; Jacob et al., 2021; Jacob & Waddell, 2020; Kaczer & Maldonado, 2009; Klappenbach, Nally, & Locatelli, 2017; McCurdy, Sareen, Davoudian, & Nitabach, 2021; Mustard, Gott, Scott, Chavarria, & Wright, 2020).

Honey bees *Apis mellifera* are used since decades as model organism to study learning and memory thanks to their remarkable abilities to form visual and olfactory memories, and to the possibility to train and test animals in restrained conditions that make them accessible for electrophysiology and calcium imaging (Giurfa & Sandoz, 2012; Rath, Galizia, & Szyszka, 2011; Strube-Bloss & Rössler, 2018). The vastly used appetitive olfactory conditioning of the proboscis extension response (PER) is based on the association between a neutral odour and a sucrose reward (Bitterman, Menzel, Fietz, & Schäfer, 1983; Takeda, 1961). After training honey bees respond extending the proboscis upon stimulation with the conditioned odour that anticipates the reward. Secondly, different aversive learning paradigms have been described in restrained honey bees too. Olfactory and gustatory stimuli can be used as conditioned stimuli to predict electric shock, heat or non-edible substances and the conditioned responses are the sting extension in some cases and suppression of the proboscis extension in others (Junca, Carcaud, Moulin, Garnery, & Sandoz, 2014; Vergoz et al., 2007; Wright et al., 2010). Experimental protocols aimed at conditioning the bees to retract their proboscis upon stimulation with an odour, require first eliciting the proboscis extension with sugar (Smith, Abramson, & Tobin, 1991) or providing a bitter substance mixed with the sugar that is used as reinforcement of the conditioned odour. In this last case, the fact that bees do not show a conditioned PER expected by the presence of sugar, is interpreted as aversive learning generated by the bitter substances that is anticipated by the odour (Wright et al., 2010). An alternative protocol for olfactory aversive learning is pairing the odour with the bitter or toxic substance alone, and assuming that an aversive association is formed even though it cannot be observed during acquisition. Whether aversive learning takes place can be observed later as a conditioned inhibition toward the odour (Ayestaran, Giurfa, & de Brito Sanchez, 2010; Rescorla, 1969). This protocol was used to evaluate the deterrent nature of different non-edible substances. However, if a stable and odour-specific memory is established remains unexplored. Here we deepen into the use of this paradigm as a tool to study olfactory aversive memory in restrained honey bees. As an advantage, the training itself does not require an appetitive context and thus training trials can be analyzed as single aversive associations. Further on, aversive training trials based on this protocol were presented intermingled with classic sucrose-rewarded trials to study appetitive and aversive memories that might be formed in parallel during the same training session. We set conditions to evaluate the independence of aversive and appetitive memories and to evidence possible interactions during acquisition and retrieval. Interestingly, in circumstances in which different memories compete, the feeding state of the animal was critical to modulate memory expression. Situations in which stimuli can predict opposite consequences represent more accurately scenes from the real world in which animals have to choose among several options the most suitable response. Here we describe an experimental design in which acquisition and retrieval of competing memories can be studied in restrained bees.

## Results

### Acquisition and duration of olfactory aversive memory

A previous study in honey bees showed that if an odour is presented paired with quinine, and shortly after that it is used as conditioned stimulus for appetitive conditioning, a retardation in the appetitive learning curve is observed (Ayestaran et al., 2010). This effect was taken as evidence of the deterrent nature of quinine and importantly, it implies an aversive association between the odour and the bitter substance. Here we took benefit of this phenomenon to study aversive learning and memory. Initially we performed the pertinent controls to validate whether pairing a neutral odour with quinine induces the formation of an associative memory. The experiment consisted of four groups of bees (figure 1 A). All of them underwent two sessions. The first session was different for each group. The bees in the first group were placed in the training position but did not receive olfactory or gustatory stimuli (placement group). The bees in the second group received odour presentations and no gustatory stimulus (odour only group). In the third group the bees were trained using an explicitly unpaired protocol. They received intermingled presentations of odour or quinine separated by 5 minutes intervals (odour–quinine group). Finally, the bees in the fourth group received odour presentation paired with quinine (odour+quinine group). Since bees did normally not extend the proboscis in response to quinine, trials with the bitter substance were accomplished by touching the antenna with a droplet of the quinine and gently extending the proboscis with a needle to touch its tip with the solution. We did not observe the animals to ingest the quinine solution, but movements that were indicative of rejecting it. The second session (testing session) was performed 2, 24 and 48 h after training. The protocol was the same for all groups and consisted in 5 presentations of the same odour used in the first session, but this time it was paired with sucrose. The existence of aversive olfactory memory should become evident as a delay or impairment in appetitive learning (Rescorla, 1969). Figure 1B shows the results obtained during the test sessions. The performance of the “odour+quinine” group was lower than the “placement” and the “odour only” groups, 2 and 24 hours after training. This result indicates that memory lasts between 24 and 48 h. The “placement” and the “odour only” groups showed similar performance which indicates that five presentations of odour alone do not induce latent inhibition toward the odour (Chandra, Wright, & Smith, 2010). An interesting result that ruled out the possibility that the low performance of the odour+quinine group was due to toxicity or aversive sensitization caused by quinine, is that the unpaired group showed facilitation of appetitive learning. This effect can be explained by a positive valence or higher salience of the odour, once the animals have learnt that the odour signals trials without the negative reinforcement (Yarali et al., 2008). Moreover, the opposite results obtained from paired and unpaired protocols, emphasises a time-discrete processing of quinine as unconditioned stimulus.

**Figure 1.**
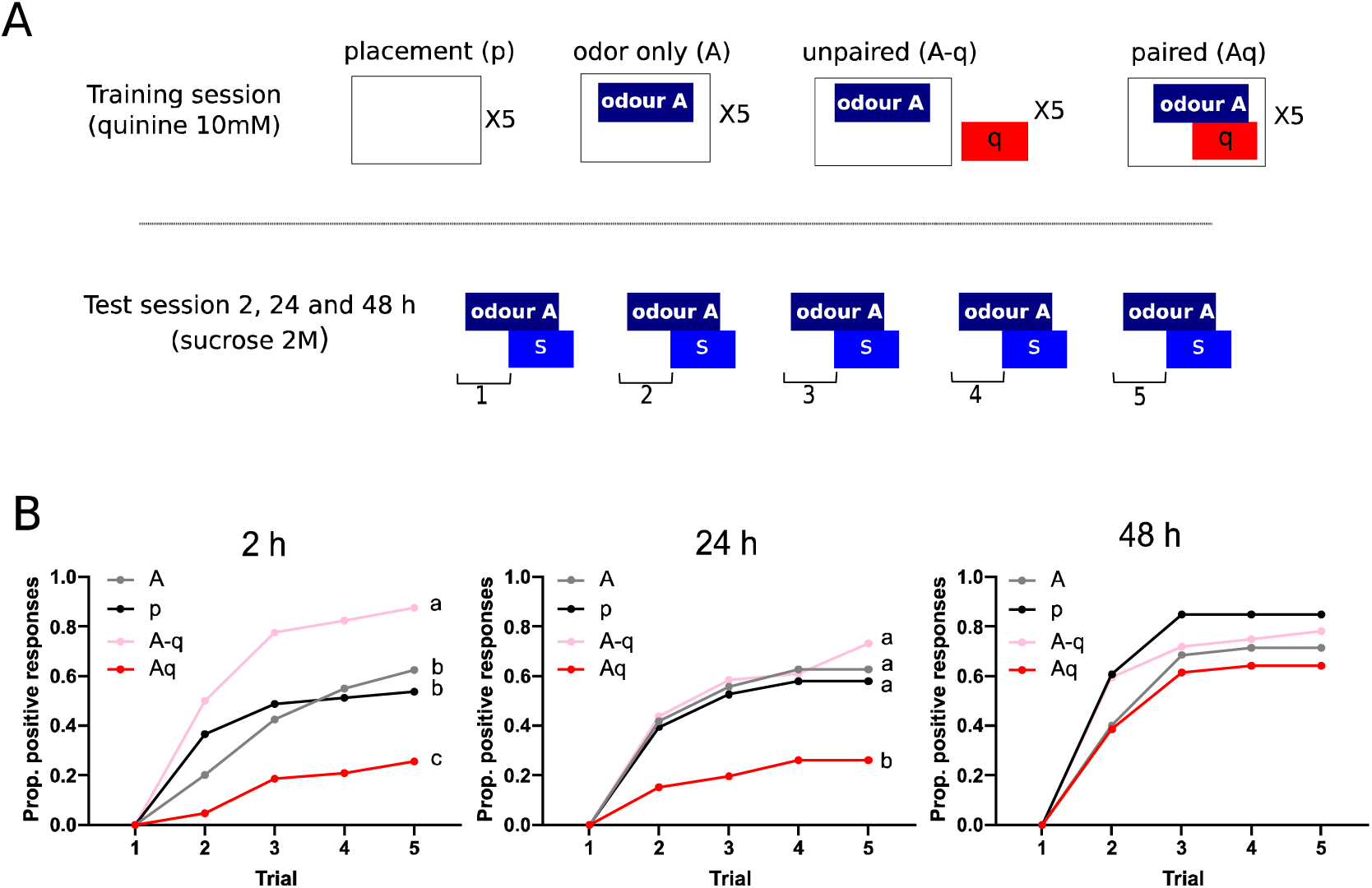
Olfactory aversive memory. **A:** Four groups of bees differed in the training session. The training session had 5 trials separated by 10 minutes intervals. Odour A was acetophenone or 1-hexanol used in balanced experiments. “q”: quinine. The test of aversive memory consisted in 5 appetitive training trials using the same odour as in the first session. “S”: sucrose reward. Test sessions were performed 2, 24 and 48 h after the first session. **B.** Proportion of bees that extended the proboscis in response to the odour and before stimulation with sucrose. Black, “placement” p group (2h: n=41; 24h: n=38; 48h: n=33); Grey, “odour only” A group (2h: n=40; 24h: n=44; 48h: n=35); Pink, unpaired “odour-quinine” A-q group (2h: n=40; 24h: n=42; 48h: n=32); Red, paired “odour+quinine” Aq group (2h: n=43; 24h: n=46; 48h: n=40). Independent groups of animals were test 2, 24 or 48 hours after the training session. **2h**: Time factor: F_3.05,489_=107, p<0.001; Group Factor: F_3,160_=15.9, p<0.001; Time x Group: F_12,640_ = 6.49; p<0.001; Contrasts: Aq vs. A, t_1,360_=5.32, p<0.001; Aq vs. P, t_1,368_=5.85, p<0.001; A-q vs. A, t_1,398_=4.83, p<0.001; A-q vs. p, t_1,402_=4.41, p<0.001; A vs p, t_1,403_=0.43, p=0.670; **24h**: Time factor: F_2.50,410_=113, p<0.001; Group Factor: F_3,164_=7.06, p<0.001 Time x Group: F_12,656_=4.12; p<0.001; Contrasts: Aq vs. A, t_1,399_=6.46, p<0.001; Aq vs. P, t_1,350_=5.53, p<0.001; A-q vs. A, t_1,417_=0.547, p=0.783; A-q vs. P, t_1,391_=1.15, p=0.582; A vs. P, t_1,398_=0.622, p=0.783; **48h**: Time factor: F_2.25,303_=194, p<0.001; Group Factor: F_3,135_=1.87, p=0.138; Time x Group: F_12,540_=1.25, p=0.247. Different letters mean p<0.05 in a Holm-Sidak post hoc test.

### Appetitive and aversive learning during the same training session

We evaluated the ability of bees to form appetitive and aversive memories acquired by intermingled positive and negative trials distributed along the same training session. The protocol constitutes a differential conditioning in which an odour is associated with a positive reinforcement and a second odour is associated with the negative reinforcement. This kind of protocols have been used before to study the ability to discriminate odours without focusing on the formation of appetitive and aversive memories (Getz, Brückner, & Smith, 1986; Getz & Smith, 1987; Wright, Choudhary, & Bentley, 2009; Wright, Kottcamp, & Thomson, 2008). The experimental design is described in Figure 2A and consisted of four groups of bees that differed in the first session. The first group received intermingled trials of two the odours (5 each) in pseudorandom sequence (odours only group, A/B). The second group received odour B paired with sucrose solution intermingled with odour A trials (appetitive trained group; A/B+). The third group received the odour A paired with quinine intermingled with odour B trials (aversive trained group, Aq/B). The fourth group received 5 trials of odour A paired with quinine intermingled with 5 trials of odour B paired with sucrose (double-trained group, Aq/B+). Figure 2B left panel, shows the learning curves of the groups that received appetitive trials (A/B+ and Aq/B+). Only a minor level of response was observed towards odour A in both groups, indicating the discriminability among these two odours and the specificity of learning. Furthermore, the appetitive learning curve toward odour B in the Aq/B+ was almost as steep as in the A/B+ group in spite of the intermingled trials with quinine. Memory retention was tested 24 hours after training. Appetitive memory was tested first with a single trial with odour B (Figure 2B, middle panel). Both groups that the day before had training protocols without appetitive trials (A/B and Aq/B) did not show proboscis extension when stimulated with odour B. In contrast, appetitive memory was clearly observed in bees of the A/B+ and Aq/B+ groups. A slight but significant impairment was observed in the double-trained group, which is consistent with the trend observed along the conditioning curve. Afterwards, all bees underwent the test session for aversive memory based on 5 appetitive conditioning trials using odour A (figure 2B right). Strikingly, no response was observed to odour A in the first test trial, which argues in favour of the animals’ ability to discriminate the two odours. The rest of the trials with odour A served to evaluate aversive memory. Both groups that had aversive training (Aq/B and Aq/B+) showed shallow learning curves in comparison to the non-aversive trained groups (A/B+ and A/B). Interestingly, aversive memory in the double trained group (Aq/B+) was also not as strong as in the only aversive trained group (Aq/B), thus, mirroring the lower performance in appetitive memory of the same group. In conclusion, the fact that Aq/B+ group shows odour specific appetitive and aversive memories, demonstrates that bees are able to form and express two memories of opposite valence acquired during the same training session.

**Figure 2.**
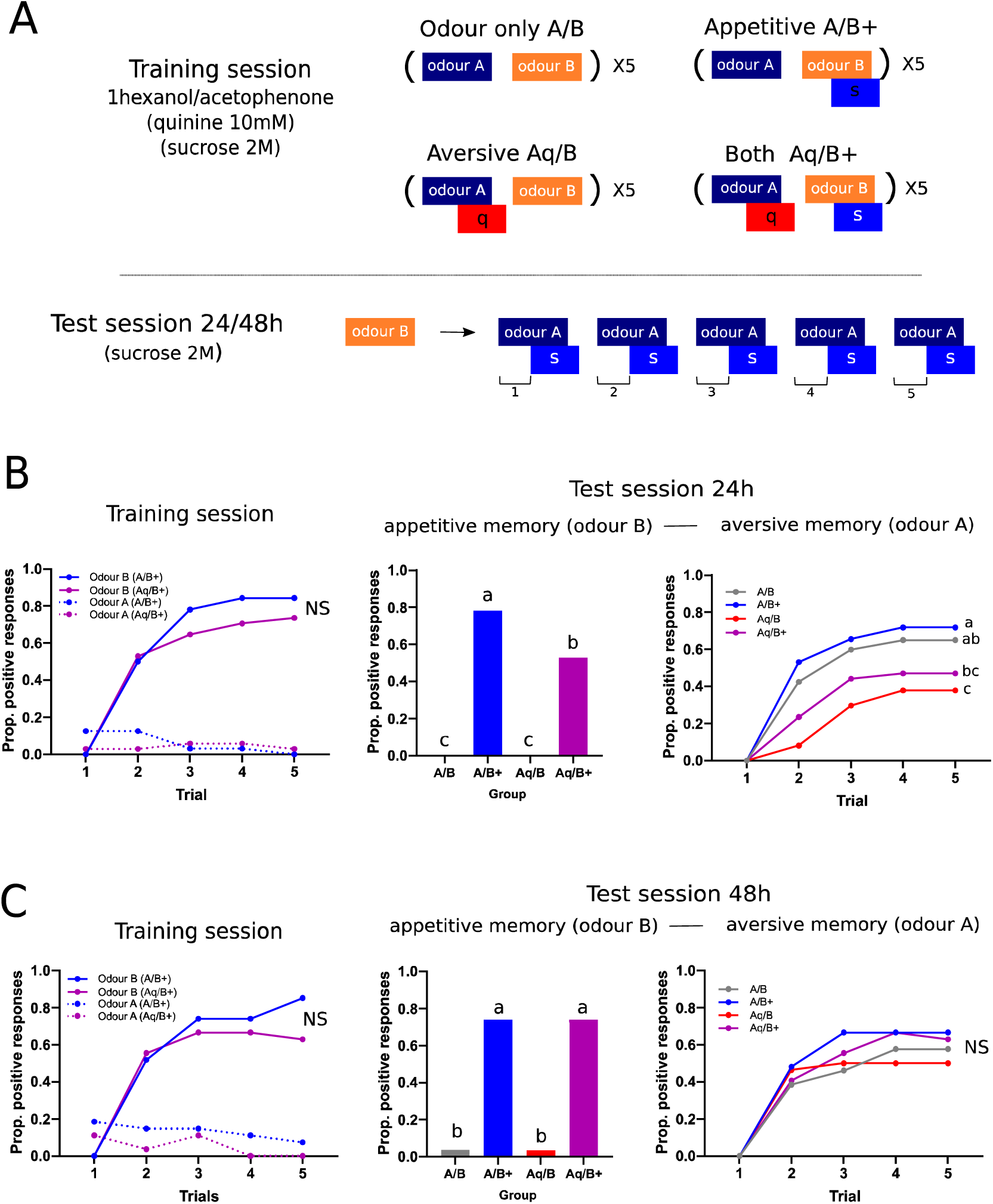
Honey bees store separate memory traces during one training session. **A.** Experimental procedure: Four groups of bees that differed in the first session. Inter-odour interval was 5 minutes. Acetophenone and 1-hexanol were used as odour A or B in counterbalanced experiments. Appetitive and aversive memory were tested 24 and 48 h after training. Appetitive memory was tested with one trial of odour B. Aversive memory was tested with 5 trials of appetitive conditioning using odour A. **B left panel,** training session. Blue, bees trained pairing odour B with sucrose and odour A without quinine (A/B+, n=32). Solid lines: responses to odour B. Dotted lines: responses to odour A. Magenta: bees that received intermingled appetitive and aversive training trials (Aq/B+, n=34). Solid lines: responses to odour B; Dotted lines: responses to odour A. Analysis of responses to odour B: Time factor: F_3.26,209_=84.0, p<0.001; Group Factor: F_1,64_=0.87, p=0.354; Time x Group: F_4,256_=1.20, p=0.312. **B middle panel,** appetitive memory test 24 h after training. Grey: animals that were exposed to both odours without sucrose or quinine (A/B, n=40). Blue: animals that received odour B paired with sucrose and odour A without quinine (A/B+, n=32). Red: animals that during the first session received odour A paired with quinine and odour B without sucrose (Aq/B, n=37). Magenta: animals that received odour B rewarded with sucrose and odour A reinforced with quinine (Aq/B+, n=34) (F_3,139_=53.62, p<0.001). Holm-Sidak Contrast: A/B+ vs Aq/B+, t _139_=3.23, p<0.01. **B right panel,** aversive memory test 24 h after training. Colours and groups same as in central panel. Time factor: F_2.20,305_=104, p<0.001; Group Factor: F_3,139_=5.43, p<0.01; Time x Group: F_12,556_=2.83; p<0.001. Holm-Sidak Contrasts: A/B vs. Aq/B: t_1,380_=5.07, p<0.001; A/B+ vs Aq/B+: t_1,323_=3.77, p<0.001; Aq/B vs Aq/B+: t_1,340_=2.04, p=0.083. **C.** Colours and groups same as in B. (A/B: n=26; A/B+: n=27; Aq/B: n=28; Aq/B+: n=27). **C left panel,** training session. Analysis of responses to odour B, Time factor: F_3.45,179_=59.3, p<0.001; Group Factor: F_1,52_=0.57, p=0.453; Time x Group: F_4,208_=1.52 p=0.198. **C middle panel,** appetitive memory test 48 h after training (F_3,100_=37.04, p<0.001). Contrast Holm-Sidak, A/B+ vs Aq/B+ t_100_=0.017, p=0.99. **C right panel:** Aversive memory test: Time factor: F_2.25,234_=91.3, p<0.001; Group Factor: F_3,104_=0.459, p=0.712; Time x Group: F_12,416_=0.91, p=0.538. Different letters mean p<0.05 in a Holm-Sidak post-hoc test.

A second set of bees was tested 48 h after the training session (Figure 2C). This time, we did not find any difference in appetitive memory in groups A/B+ and Aq/B+ (Fig 2C mid panel). Moreover, no difference was observed among groups along the test session for aversive memory. This lack of aversive memory in the Aq/B and Aq/B+ groups is consistent with the results obtained in the first section that circumscribed the duration of this aversive memory to less than 48 h. Based on this last result, we interpret that the slight impairment observed in appetitive memory 24h after double training, can be explained as an interference during retrieval rather than as interference during acquisition or memory storage. This conclusion is indicated by the fact that expression of appetitive memory in the double trained group was restored to the level observed in the A/B+ group once aversive memory has vanished.

### Aversive odour embedded in a mixture

Honey bees are able to detect appetitive conditioned odours embedded in complex mixtures (Locatelli et al., 2013; Reinhard, Sinclair, Srinivasan, & Claudianos, 2010). Here, we evaluated whether honey bees are able to detect aversive learned odours embedded in binary mixtures. We trained animals using the aversive conditioning protocol and evaluated aversive memory using a binary mixture that contained the learned and a novel odour. Figure 3A shows the experimental procedure. One group was exposed to 5 presentations of odour with no unconditioned stimulus (A) and a second group received 5 trials of the odour paired with quinine (Aq). Figure 3B shows the learning curves during a test session performed 24 h later. The lower performance of the Aq group is consistent with aversive memory. Figure 3C shows a test session performed 48 h after training. As it could be expected based on the duration of this memory, performance was not different among groups at this time.

**Figure 3.**
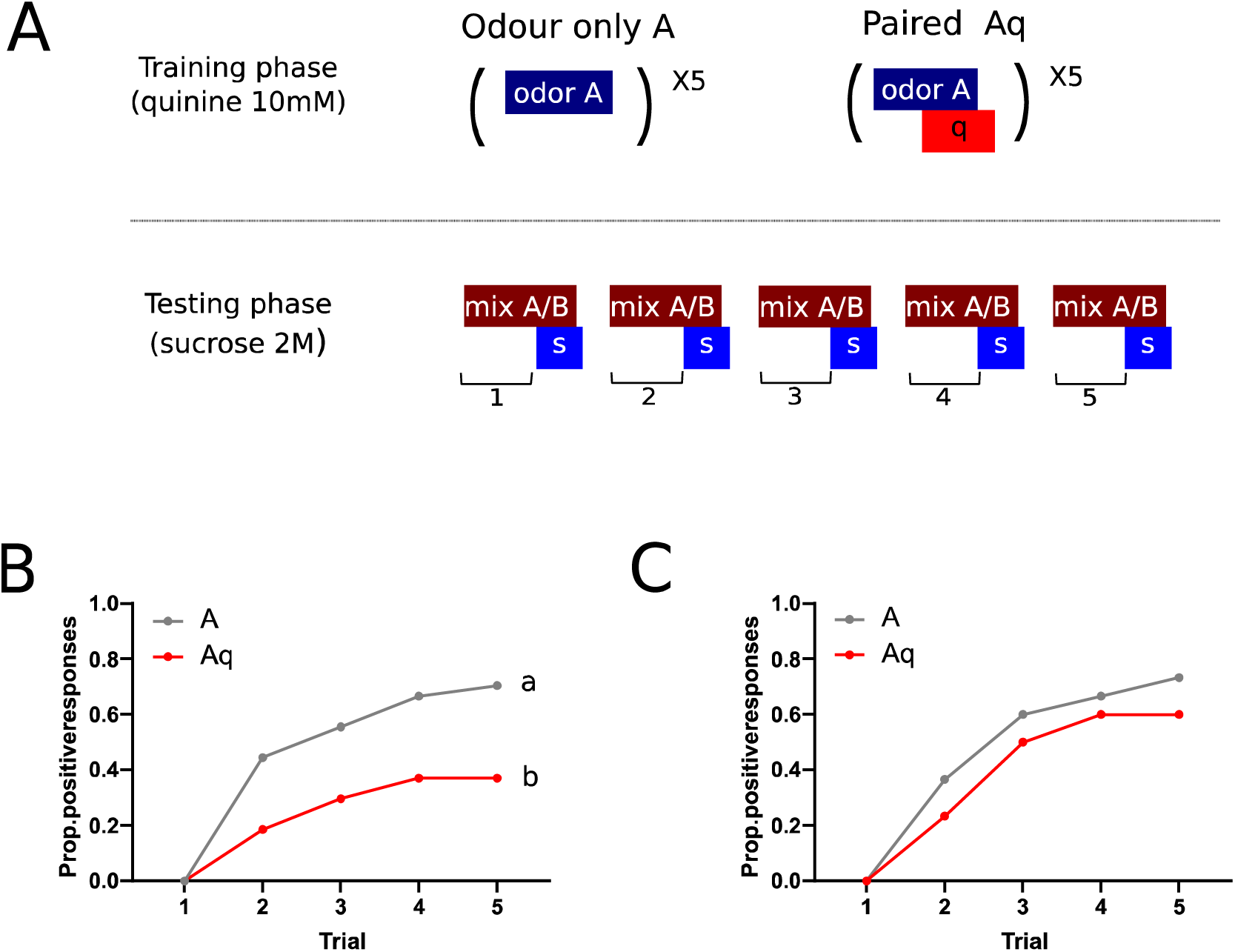
Honey bees detect aversive learned odours embedded in binary mixtures. **A.** Scheme of the experimental procedure and two groups of bees that differed in the first session. Acetophenone and 1-hexanol were used as novel and learned odours in counterbalanced experiments. Aversive memory was tested using a binary mixture (actophenone : 1-hexanol). **B.** Aversive memory test 24 h after training. Grey: animals that were exposed to the odour without quinine (A, n=45); Red: animals that received paired presentation of odour and quinine (Aq, n=43). Trial factor: F_2.53,132_=33.2, p<0.001; Group Factor: F_1,52_=5.97, p<0.05; Trial x Group: F_4,208_=3.01, p<0.05. Different letters mean p<0.05 in a Holm-Sidak post-hoc test. **C.** Aversive memory test 48 h after training. Colours and groups same as in B. (A, n=30; Aq, n=30). Trial factor: F_2.64,153_=52, p<0.001; Group Factor: F_1,58_=0.991, p=0.324; Trial x Group: F_4,232_=0.52, p=0.724.

### Appetitive and aversive memories compete during retrieval

We addressed how do honey bees perceive a binary mixture that contains appetitive and aversive learned odours. Figure 4A depicts the training and testing protocol. The training session consisted of the same four groups as in experiment 2: A/B; A/B+; Aq/B and Aq/B+. Figure 4B left panel shows the training curves of the two groups that had appetitive conditioning trials during training. Both groups (A/B+ and Aq/B+) show steep learning curves towards odour B, and no or only minimal response towards odour A as observed in experiment 2. During the second session (figure 4B right) all bees were tested using a binary mixture of the odours A and B, each odour in the same concentration used during training. The untrained group (A/B) shows a standard acquisition curve, i.e. no bee responded to the odour during the first trial and 60% of them responded in the fifth trial. The Aq/B group showed a shallow acquisition curve which is consistent with aversive memory and the ability to detect the presence of odour A in the mixture. The A/B+ group showed high response from the first trial of the test session, consistent with appetitive memory and the ability to recognize the presence of the odour B in the mixture. Surprisingly, the performance of the Aq/B+ group did not differ from the A/B+ group (blue and magenta) behaving as only expressing appetitive memory.

**Figure 4.**
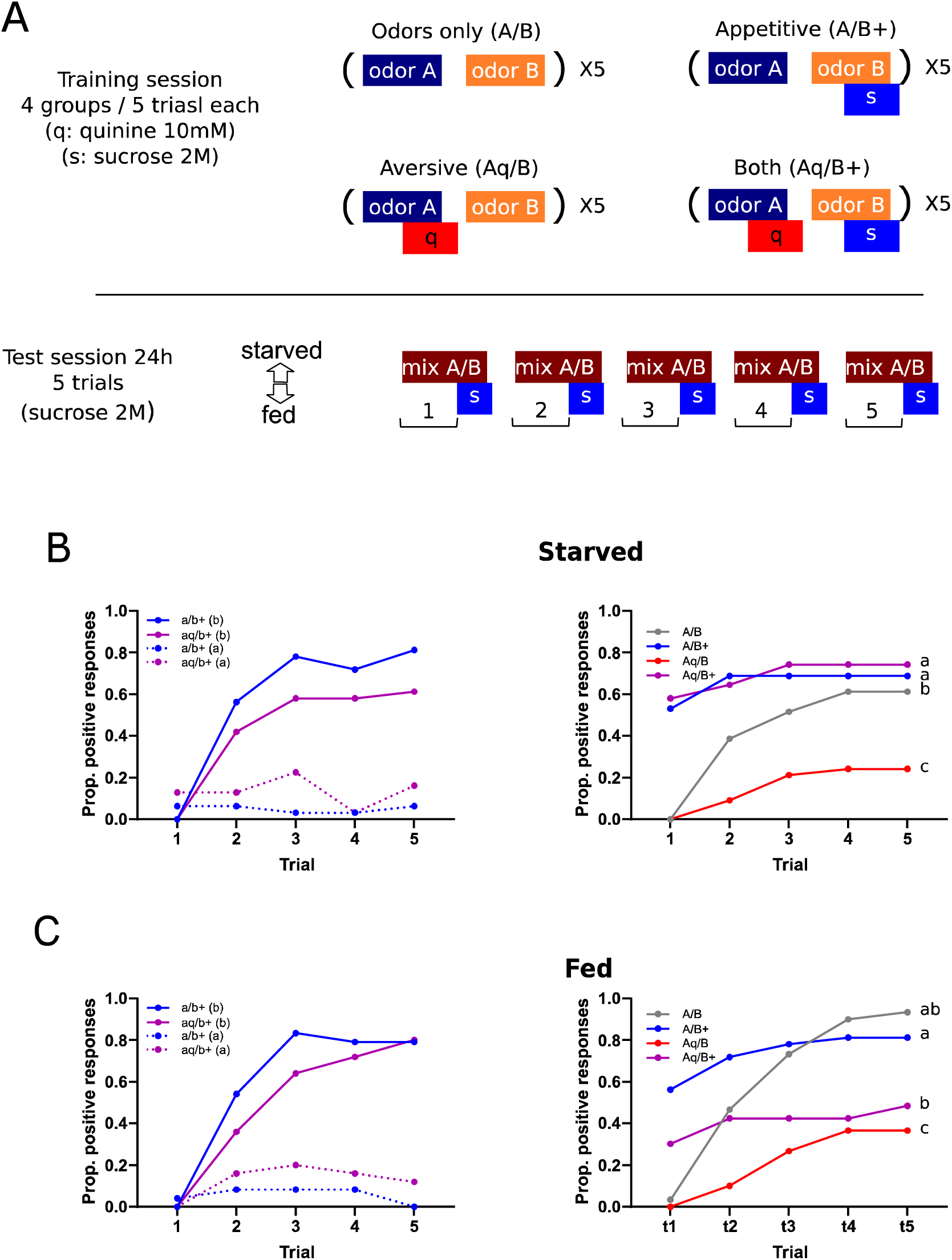
Honey bees can switch from expressing appetitive to aversive memory. **A.** Experimental procedure and four groups of bees that differed in the first session. Memory was tested using a mixture of odours A and B. Acetophenone and 1-hexanol were used as A and B in counterbalanced experiments. **B left**, acquisition curves during training. Blue: animals that were trained pairing odour B with sucrose and odour A without quinine (A/B+, n=32). Solid lines: proportion of bees that responded to odour B. Dotted lines: proportion of bees that responded to odour A. Magenta: bees that received intermingled appetitive and aversive trials (Aq/B+, n=31). Solid lines: responses to odour B; Dotted lines: responses to odour A. Analysis of response to odour B, Time factor: F_3.29,201_=59.5, p<0.001 ; Group Factor: F_1,61_=2.78, p=0.101; Time x Group: F_4,244_=1.14, p=0.340. **B right**, acquisition curves during test. Feeding conditions same as in previous experiments (i.e. 18 to 22 h since last feeding). Grey: animals that in the first session were exposed to both odours without sucrose or quinine (A/B, n=31). Blue: animals that during the first session received odour B paired with sucrose and odour A without quinine (A/B+, n=32). Red: animals that during the first session received odour A paired with quinine and odour B without sucrose (Aq/B, n=33). Magenta: animals that during the first session received odour B rewarded with sucrose and odour A reinforced with quinine (Aq/B+, n=31). Time factor: F_1.98,244_=39.1, p<0.001; Group Factor: F_3,123_=12.9, p<0.001; Time x Group: F_12,492_=5.32, p<0.001. Holm-Sidak Contrasts: A/B vs. Aq/B: t_1,235_=5.90, p<0.001 (aversive memory); A/B+ vs. Aq/B+: t_1,250_=0.52, p=0.601 (appetitive memory). **C,** Groups and colours same as in B, except that bees were fed 2 h before the second session. (A/B: n=30; A/B+: n=32; Aq/B: n=31; Aq/B+: n=34). **Left panel,** Time factor: F_3.08,145_=56.5, p<0.001; Group Factor: F_1,47_=1.14, p=0.292; Time x Group: F_4,188_=1.17, p=0.327. **Right panel**, Time factor: F_3.49,328_=38.4, p<0.001; Group Factor: F_3,94_=32.8, p<0.001; Time x Group: F_12,376_=9.32, p<0.001. Contrasts: A/B vs. Aq/B: t_1,237_=6.10, p<0.001 (aversive memory); A/B+ vs Aq/B+: t_1,252_=6.02, p<0.001 (aversive memory). Different letters mean p<0.05 in a Holm-Sidak post hoc test.

We hypothesized that expression of appetitive memory might be occluding aversive memory, and if so, the expression of the last one could be restored if the weight of the appetitive memory in the decision weren’t so high. Therefore, we repeated the experiment, but we fed the animals 2 hours before test (1.5 μl of 1M sucrose solution) in order to reduce appetite. Figure 4C right panel corresponds to the test session. Bees in the A/B group (grey) showed a regular acquisition curve indicating that the amount feeding did not affect appetitive learning. In the A/B+ group (blue), 60% of the bees started responding from first trial of the test session, a response level that is consistent with appetitive memory. The performance of these two groups is important, since they show that feeding did not suppress appetitive behaviour. Interestingly, this time the performance of the Aq/B+ group (magenta) was significantly lower than the A/B+ group (blue). This change is due to a number of bees that would have expressed appetitive memory, but because they were fed, they switched from expressing appetitive to expressing aversive memory. This interpretation is further supported by the fact that the bees in this group didn’t almost changes their decision along the whole test session, at contrast with what happened in the A/B group. In summary, feeding the animals unveiled aversive memory and provided evidence that appetitive and aversive memory traces are ready to be expressed depending on the internal state.

### Appetitive and aversive memories in the same individuals

In the second and fourth sections we evaluated the behaviour of bees that underwent double training protocols (Aq/B+). We concluded that honey bees are able to form appetitive and aversive memories acquired during the same training session. This interpretation was, however, based on population’s behaviour. It might happen that a fraction of bees expressed aversive memory and a non-overlapping fraction expressed appetitive memory. Therefore, we re-analysed the data based on individual performance. In experiment 2, animals were first tested for appetitive memory using a single trial with odour B. Bees that extended the proboscis to odour B were considered as expressing appetitive memory. Immediately after that, the bees were tested with five appetitive conditioning trials using odour A. We considered that the bees that did not respond in any of the five test trials with odour A were affected by aversive memory. Thus, we classified the Aq/B+ bees (n=34) in four categories: no memory (12 %), only appetitive memory (35%), only aversive memory (35%) and both memories (18%). Rephrasing it, 33% of bees that showed appetitive memory did also show aversive memory, and 33% of bees that showed aversive memory did also show appetitive memory. This indicates that both memories are not mutually exclusive. However, we performed a test for independence among appetitive and aversive memories, and found that the number of bees showing both memories was significantly lower than expected if the two classifications were independent (Chi2 df=1, p=0.015). Thus, even though it is possible for honey bees to store and express both memories, there is also some degree of interference. This conclusion coincides with the observation in experiment 2 that appetitive and aversive memories in the double-trained group were reduced in comparison to memory measured in the single trained groups (Figure 2). Interestingly, when the bees tested 48 h after training were analyzed in the same way, the proportion of bees that did not respond to odour A in the test session dropped from 52% 24h after training to 33% 48h after training, while the animals expressing appetitive memory increased from 52% to 74%. Thus, the fact that the number of bees expressing appetitive memory increases concomitantly with a reduction in the number of bees expressing aversive memory suggests that the interference measured 24h after training occurred during memory retrieval, rather than during its acquisition or consolidation. In experiment 4 we cannot easily determine if an animal has both memories, because the ways to express them are mutually exclusive. However, we quantified the bees that responded extending the proboscis in the first test trial as bees expressing appetitive memory, and the bees that during the test session did not respond in any of the five test trials with odour A as bees expressing aversive memory. This analysis shows that the proportion bees showing appetitive memory in the double-trained group dropped from 58% before feeding to 36% after feeding, while the bees showing aversive memory increased from 26% before feeding to 48% after feeding, thus, providing evidence that aversive memory was present in the double-trained bees but occluded by the expression of appetitive memory.

## Discussion

### Reversal learning as a tool for revealing aversive memory

In the classic appetitive olfactory conditioning of the PER in honey bees, a neutral odour is paired with sucrose solution applied to the antennae and proboscis. Once the association is established, the sole stimulation with the conditioned odour causes extension of the proboscis, a fact that is taken as evidence of appetitive memory (Bitterman et al., 1983; Takeda, 1961). On the other side, studies conceived to show learned suppression of the proboscis extension were based on eliciting its extension with sugar and signalling the occurrence of an electric shock or quinine with an odour, otherwise active retraction of the proboscis cannot be monitored (Smith et al., 1991; Wright et al., 2010). Here we used a different strategy which does not need stimulation with sucrose during aversive training. A neutral odour is presented paired with quinine. No evidence of learning can be measured during the training session. Whether a memory was built can be measured later, during a test session in which the same odour is paired with sucrose solution. This second session constitutes a reversal learning protocol in which animals have to assign the odour a new value that is opposite to the one learned before and thus, the appetitive learning curve is severely affected (Devaud, Blunk, Podufall, Giurfa, & Grünewald, 2007; Hadar & Menzel, 2010). In the study by Ayestaran et al. (2010) this phenomenon was used to evaluate the deterrent nature of quinine and other bitter substances. In the present work, we use it to study learning and memory, and determined that a training protocol of five spaced trials of a neutral odour paired with quinine induces the formation of a memory that lasts between 24 and 48 h. Based on the observation that the bees did not ingest the quinine solution during training trials, we conclude that in the present case aversive reinforcement relies on the gustatory modality upon touching receptors on the proboscis. However, since we cannot discard that a minimal amount of quinine solution might have been ingested, pre and post-ingestive pathways might have signalled the negative reinforcement (Wright et al., 2010).

Interestingly, explicitly unpaired presentations of odour and quinine facilitated subsequent appetitive learning of the same odour (figure 1B 2h). This result is important two-fold. First, it rules out that the retardation in appetitive learning after a treatment with quinine is due to toxicity or malaise (Hurst, Stevenson, & Wright, 2014). If this were the case, quinine should affect appetitive learning regardless if it was applied paired or unpaired. Second, it reinforces the conclusion that this training protocol induces associative learning and not a quinine-induced aversive sensitization that produces unspecific suppression of the proboscis extension. Interestingly, the consequence of unpaired CS-US presentations resembles the observation reported in the seminal work by Bitterman et al. (1983), in which it was shown that unpaired stimulations with odour and sugar produced a conditioned inhibition that affected subsequent appetitive learning of the same odour. Furthermore, the possibility to convert an aversive learning protocol into an appetitive one by altering the timing between the CS and an aversive US has been studied and compared across humans, rats and flies (Andreatta et al., 2012). In flies, presenting a neutral odour shortly after a negative reinforcement, provides this odour a positive valence, likely because as in relief learning the odour signals that the shock has finished (Aso & Rubin, 2016; König et al., 2018; Vogt, Yarali, & Tanimoto, 2015; Yarali & Gerber, 2010). Furthermore, in the context of differential aversive conditioning, it turned out that in addition to aversive memory towards the paired odour, flies establish a complementary safety memory that is expressed as preference toward the control odour that during training is presented unpaired to the shock (Jacob & Waddell, 2020).

The expression of the memory that we describe here is akin to latent inhibition, i.e. an odour specific retardation of acquisition in a subsequent appetitive conditioning (Chandra et al., 2010). This similarity puts the question whether latent inhibition might actually constitute a sort of aversive learning, as supported by studies in flies (Jacob et al., 2021), or if paired exposures to odour and quinine does actually accelerates latent inhibition which normally requires 20 trials or more. Previous studies in honey bees ruled out the first interpretation by showing that unrewarded odour exposures that produce latent inhibition do not convert the odour into a conditioned inhibitor (Chandra et al., 2010; Fernández, Giurfa, Devaud, & Farina, 2012). In regards to the second possibility, we obtained different results depending on the way in which memory was evaluated. In experiment 1 the animals were tested with the odour that was used for aversive training, and the performance is compatible with both, aversive memory and latent inhibition. In experiment 3, the odour that had been paired with quinine was presented during test together with a novel odour. The observed inhibition in appetitive learning is consistent with a negative value of the odour rather than with ignoring it. Finally, in experiment 4, the aversive learned odour was presented during test together with an appetitive learned odour. This time, honey bees behaved as ignoring or as avoiding the aversive learned odour depending on their satiation level, which would be consistent with latent inhibition in the first case and with aversive memory in the second. These results prompt further mechanistic studies to understand whether the ability to behave as ignoring an aversive stimulus is the result of reducing its negative value or accepting its inherent risk.

### Two memories coexist after differential conditioning

One of our main objectives was to study the ability of bees to extract and use information from experiences that mix appetitive and aversive associations. Taking benefit of the aversive conditioning protocol described in the first section, we addressed whether honey bees are able to form appetitive and aversive memories during a training session that intermingles appetitive and aversive training trials. A previous study provided evidence that honey bees can learn that a given odour predicts an electric shock and a different one predicts sugar, both presented in the same training session (Vergoz et al., 2007). Here we have challenged the bees to an even more difficult task, since both appetitive and aversive unconditioned stimuli involve the gustatory modality and, furthermore, the expression of both memories require extending the proboscis in one case and suppressing the extension in the other. Interestingly, bees in the double-trained group behaved as having formed both memories without mixing the learned value of each odour. Several observations lead us to conclude that appetitive and aversive memories are independently acquired and that they compete during expression. First, we observed only a slight but not significant reduction of the appetitive learning curve during the training session of the double-trained groups. Second, during memory test 24 h after training, aversive and appetitive memories in the double-trained group were significantly reduced compared to simple-trained groups. Up to that point, a possible interpretation could be a mutual interference during memory consolidation. However, we observed in experiment 2, that appetitive memory was fully recovered when aversive memory vanishes 48 h after training. A similar effect was observed in experiment 4, in which expression of aversive memory was recovered when honey bees were partially fed to reduce appetitive arousal. Thus, the fact that expression of appetitive or aversive memories is restored when the opposite one reduces its expression indicates that both memory traces are established after double training, and that expression of one of them can occlude the expression of the other. Then, if the reduction in memory expression is caused by the fact that both memories cannot be expressed at the same time, why is this effect also observed when the tests with odours A and B were split (experiment 2; 24h test)? At this point, a certain weight must be conceded to the training context as part of the conditioned stimulus. Appetitive and aversive trainings take place in the same visual and mechanosensory context that may also become predictors of sugar and quinine. It is reasonable to expect that the expression of olfactory memory competes with context memory and affect performance (Gerber & Menzel, 2000).

The fact that that honey bees in the double trained group managed to establish the specific predictive value of each odour without mixing this information highlights the independence of the appetitive and aversive pathways involved in learning and memory formation. A number of studies based on different strategies and using different learning paradigms in honey bees provided consistent proof that the biogenic amine octopamine provides the internal signal necessary for appetitive learning, while dopamine and serotonin are necessary for aversive learning induced by quinine and electric shocks (Farooqui et al., 2003; Hammer & Menzel, 1998; Lai et al., 2020; Vergoz et al., 2007; Wright et al., 2010). It must also be considered that the time interval between appetitive and aversive training trials might have contributed to the CS-US specificity and the lack of interference during acquisition. Based on previous pharmacological studies, it is expected that if appetitive and aversive unconditioned stimuli occur simultaneously or close in time, interference should be expected during acquisition. Indeed, it has been shown that administration of octopamine during aversive learning affects aversive memory formation (Agarwal et al., 2011) and administration of dopamine during appetitive learning affects appetitive memory (Klappenbach, Kaczer, & Locatelli, 2013).

### Aversive learned odours embedded in mixtures

Honey bees are able to find learned odours embedded in complex mixtures (Chen et al., 2015; Reinhard et al., 2010). However, this ability was tested only for appetitive learned odours. Here we provide evidence that honey bees can also react to aversive learned odours embedded in mixtures. This ability is particularly relevant in nature, where meaningful odours are immersed in changing and noisy backgrounds (Conchou et al., 2019; Raguso, 2008). The mechanisms by which honey bees, and other animals, detect the presence of key odourants embedded in complex mixture are still matter of intense research (Marachlian, Klappenbach, & Locatelli, 2021). Behavioural studies have shown that odour mixtures can be perceived by honey bees in elemental and configural ways (Deisig, Lachnit, & Giurfa, 2002; Deisig, Lachnit, Sandoz, Lober, & Giurfa, 2003; Reinhard et al., 2010; Smith, 1998). Physiological studies have also found elemental and configural representations of odour mixtures depending on where it is measured along the olfactory circuit (Deisig, Giurfa, Lachnit, & Sandoz, 2006; Deisig, Giurfa, & Sandoz, 2010; Krofczik, Menzel, & Nawrot, 2009; Yamagata, 2009). Our results based on generalization after appetitive and aversive conditioning support the view that honey bees are able to detect and recognize the presence of learned elements in a binary mixture. These results do not discard that the mixture may also produce a unique or configural perception, while still preserving information about the single components (Deisig et al., 2003; Lei & Vickers, 2008). Interestingly, the fact that honeybees can detect a learned odour embedded in a mixture correlates with experience-dependent changes in the representation of the mixtures in the AL (Marachlian et al., 2021). We have already described that mixture representation at the level of projection neurons changes depending on the experience with the pure components. The ensemble of projection neurons that encode a mixture is more similar to the ensemble of neurons that encodes rewarded components than the non-rewarded ones (Chen et al., 2015). Moreover, when honey bees learn to ignore an odour after a latent inhibition protocol, a reduction is observed in the contribution of that odour to the representation of a mixture (Andrione, Timberlake, Vallortigara, Antolini, & Haase, 2017; Locatelli et al., 2013). Future studies will have to address if aversive learning does also produce a change in the representation of a mixture that contains aversive learned odours. Interestingly, the results obtained in experiment 4 with starved and fed bees, suggest that the relative weight that each component has on the perceptual quality of a mixture might not be only determined by previous experience, rather it might be also tuned by the physiological state of the animal.

### Opposite memory traces compete for expression

In the last experiment we evaluated which one is the valence that honey bees assign to a stimulus that predicts appetitive and aversive consequences. We took animals that had undergone double training and tested them with a mixture of the appetitive and aversive learned odours. In this experiment only a 16% of bees showed no memory. The rest of them behaved as expressing appetitive or aversive memory. Thus, we conclude that mixing appetitive and aversive learned odours does not convert the mixture into a neutral stimulus, which could happen if the memory traces compete in a way in which they cancel each other. Most of the bees behaved as if they were reacting to the presence of the appetitive learned odour, however, if they were partially fed before the test session, a fraction of them behaved as detecting the presence of the aversive learned odour. Importantly, the satiation level after partial feeding did not affect the expression of the appetitive memory in the appetitive trained group (A/B+) or the ability of the control group (A/B) to learn during the test session. Decision-making in real-life situations must take into account appetitive and aversive consequences assessed in the context of the individual’s needs. Food is rewarding in many circumstances, but it can be rejected if it involves unintended consequences, or accepted if there is no other option (Desmedt, Hotier, Giurfa, Velarde, & De Brito Sanchez, 2016). Here we have assessed this situation in relation to learning and memory. We have set experimental conditions that intend to replicate a realistic situation in which appetitive and aversive memory traces compete under controlled conditions. All results are consistent with a model in which both memories are independently stored and can be alternatively expressed depending on the animal’s requirements at the moment of retrieval. Interestingly, we reached similar conclusions in a completely different animal model and behavioural tasks (Klappenbach et al., 2017). Crabs of the species *Neohelice granulata* were double-trained in a fear conditioning and an appetitive conditioning using the same context as CS. When the crabs were put back in the context they expressed either appetitive or aversive memory depending on whether they had been fed or not before the test. Results obtained in flies *Drosophila melanogaster* support the same conclusion that aversive and appetitive experiences linked to the same conditioned stimulus are stored and retrieved as independent memories (Das et al., 2014; Felsenberg et al., 2018; Perisse et al., 2013). Studies in Drosophila pointed out that the ability of a conditioned stimulus to switch from evoking appetitive or aversive responses is provided by the balance of inputs that confer information about the internal state at the output lobes of the mushroom bodies(Senapati et al., 2019).

Neural changes that accompany memory formation in honey bees have been mapped all the way from the sensory neurons and antennal lobes, to the mushroom bodies calyces and output lobes (Chen et al., 2015; Claudianos et al., 2014; Locatelli, Fernandez, & Smith, 2016; Locatelli et al., 2013; Rath et al., 2011; Strube-Bloss, Nawrot, & Menzel, 2011). Future studies will have to address how long-lasting neural changes related with the predictive value of a given odour, and transient changes that readjust the weight of these memories during retrieval, interact along these circuits to ensure adaptive behaviour. Here we provide a robust behavioural approach that can be used to induce and measure these phenomena in restrained bees.

## Materials and methods

### Animals

Honey bee (*Apis mellifera*) pollen-foragers were collected at the entrance of two regular hives situated at the Campus of the University of Buenos Aires. Bees were captured in glass vials and immobilized by shortly cooling them on ice and restrained in individual holders. Dental wax was used to fix animal’s heads in a way that they could move antennae and proboscis. After recovery from cooling, bees were fed 5 µl of 1.0 M sucrose solution and remained undisturbed until the evening when they were fed *ad libitum*. At the laboratory, bees were kept in a humid box at room temperature (20–24 ºC) on a 12:12 h light:dark cycle. All training and testing sessions were carried out between 10:00 AM and 1:00 PM. Thirty minutes before training all animals passed an admission test that consisted in touching the antennae with 2.0 M sucrose solution. Only animals that showed a rapid and conspicuous extension of the proboscis were used in the experiments.

### Odour stimulation

Odours used were 1-hexanol and acetophenone diluted 1/10 in mineral oil (all reagents from Sigma-Aldrich). Both odours were used in all experiments in counterbalanced way. One hundred µl of the odour dilution was fresh loaded into sealed 5 ml glass vials before the experiments. The odour delivery device provided a continuous 500 ml /min stream of charcoal-filtered air pointed toward the bee’s head. During odour stimulation, a solenoid valve was activated to derive part of the air flow (50ml/min) to the vial containing the odourant and a fraction of the head-space inside the vial was pushed into the main airstream in a mixing chamber before reaching the animal. Odour mixtures were obtained by activating the airflow through two parallel vials systems that converged in the mixing chamber. The whole system was designed and controlled to provide the same final volume and speed in the air reaching the bees, thus no change in total air-flow was produced during onset or offset of odour stimulation. When a mixture was used, the final concentration of each odour was the same as when it was used alone. Odour stimulation lasted always 4 seconds. Odours were removed from the training arena by a continuous and gentle exhaust placed 10 cm behind the honey bee.

### Appetitive training

Honey bees were trained using olfactory conditioning of the proboscis extension reflex (PER) (Bitterman, Menzel, Fietz, & Schäfer, 1983; Takeda, 1961). Appetitive conditioning was used to test aversive memory and also during training in experiments 2 and 4. In all cases, each appetitive conditioning trial followed the same design. Odour was used as the conditioned stimulus and 2M sucrose solution as unconditioned stimulus. In each training trial an animal was individually positioned in the training arena facing toward the odour delivery device that provided a constant stream of filtered air. The bee remained undisturbed in this position during 20 seconds and then the odour started and lasted 4 sec. Three seconds after odour onset the antennae were touched with a 2M sucrose-solution which elicits proboscis extension. Sucrose was manipulated and offered to the bee using a glass Gilmont GS-1200 Micrometer Syringe and a metal needle. When the proboscis was extended the bee was allowed to lick a droplet of 0.4µl of solution. Ingestion of this amount of solution did never take longer than 4 seconds. Twenty seconds after the end of the reward the bee was returned to the rest position apart of the training arena until the next trial.

We used in all cases a training protocol that consisted of 5 trials. The test sessions for appetitive memory in experiment 2 consisted of one 4s of odour presentation without reward. The response of each subject was recorded as positive if the subject extended its proboscis beyond a virtual line between the open mandibles during the stimulation with the odour and before stimulation with sucrose. Proportion of response was calculated for each trial as the number of bees that extend the proboscis over the total number of bees.

### Aversive training

We used an olfactory conditioning protocol based on quinine as aversive gustatory unconditioned stimulus (Ayestaran et al., 2010; Wright et al., 2010). Bees were placed in the same training position as used for appetitive conditioning protocol. Three seconds after odour onset the antennae were touched with droplet of a 10mM quinine (Quinine hydrochloride, Sigma-Aldrich Q1125) prepared in distilled water. As this normally does not elicit proboscis extension, we gently forced its extension with a needle and touched the proboscis with the solution during 4 seconds. Same as with sucrose, quinine solution was manipulated and offered using a micrometer syringe with a metal needle. Volume was indistinct since bees did not ingest it. Twenty seconds after the end of this stimulation the bee was returned to the rest position until the next trial. The training protocol consisted of 5 trials separated by 10 min intervals. The test for aversive memory consisted in 5 trials of the standard appetitive conditioning protocol described the previous section (Appetitive training).

### Statistical analysis

With the exception of the appetitive test showed in figure 2 (One way ANOVA and Holm-Sídák multiple comparisons) data were analyzed by repeated-measures General Linear Models (RM-GLM), using experimental groups and trials as fixed factors. When appropriate, Holm-Sídák multiple comparisons test between groups was performed (GraphPad Prism 8). Figures were designed with Inkscape™.

## Acknowledgments

The study was supported by ANPCyT, Ministry of Sciences, Argentina: PICT2017-2284 to FFL and PICT2017-1285 to MK ; and Universidad de Buenos Aires: UBACyT: 20020170100736BA to FFL. MK, AEL and FFL are suportad by Consejo Nacional de Investigaciones Científicas Técnicas (CONICET) Argentina.

